# An immunoevasive strategy through clinically-relevant pan-cancer genomic and transcriptomic alterations of JAK-STAT signaling components

**DOI:** 10.1101/576645

**Authors:** Wai Hoong Chang, Alvina G. Lai

## Abstract

Since its discovery almost three decades ago, the Janus kinase (JAK)-signal transducer and activator of transcription (STAT) pathway has paved the road for understanding inflammatory and immunity processes related to a wide range of human pathologies including cancer. Several studies have demonstrated the importance of JAK-STAT pathway components in regulating tumor initiation and metastatic progression, yet, the extent of how genetic alterations influence patient outcome is far from being understood. Focusing on 133 genes involved in JAK-STAT signaling, we found that copy number alterations underpin transcriptional dysregulation that differs within and between cancer types. Integrated analyses on over 18,000 tumors representing 21 cancer types revealed a core set of 28 JAK-STAT pathway genes that correlated with survival outcomes in brain, renal, lung and endometrial cancers. High JAK-STAT scores were associated with increased mortality rates in brain and renal cancers, but not in lung and endometrial cancers where hyperactive JAK-STAT signaling is a positive prognostic factor. Patients with aberrant JAK-STAT signaling demonstrated pan-cancer molecular features associated with misexpression of genes in other oncogenic pathways (Wnt, MAPK, TGF-β, PPAR and VEGF). Brain and renal tumors with hyperactive JAK-STAT signaling had increased regulatory T cell gene (Treg) expression. A combined model uniting JAK-STAT and Tregs allowed further delineation of risk groups where patients with high JAK-STAT and Treg scores consistently performed the worst. Providing a pan-cancer perspective of clinically-relevant JAK-STAT alterations, this study could serve as a framework for future research investigating anti-tumor immunity using combination therapy involving JAK-STAT and immune checkpoint inhibitors.

## Introduction

In their quest to survive and prosper, tumor cells are armored with a unique ability to manipulate the host’s immune system and promote pro-inflammatory pathways. Inflammation can both initiate and stimulate cancer progression, and in turn, tumor cells can create an inflammatory microenvironment to sustain their growth further(1, 2). Cytokines are secretable molecules that influence immune and inflammatory processes of nearby and distant cells. Although cytokines are responsible for inflammation in cancer, spontaneous eradication of tumors by endogenous immune processes rarely occurs. Moreover, the dynamic interaction between tumor cells and host immunity shields tumors from immunological ablation, which overall limits the efficacy of immunotherapy in the clinic.

Cytokines can be pro-or anti-inflammatory and are inter-dependent on each other’s function to maintain immune homeostasis(3). Discovered as a critical regulator of cytokine signaling, the Janus kinase (JAK) - signal transducer and activator of transcription (STAT) pathway allows cytokines to transduce extracellular signals into the nucleus to regulate gene expression implicated in a myriad of developmental processes including cellular growth, differentiation and host defense(4). JAK proteins interact with cytokine receptors to phosphorylate signaling substrates including STATs. Unlike normal cells which phosphorylate STATs temporarily, several STAT proteins were found to be persistently phosphorylated and activated in cancer(5). Studies have shown that persistent activation of STAT3 and STAT5 promote inflammation of the microenvironment, tumor proliferation, invasion and suppress anti-tumor immunity(5). Particularly in cancers associated with chronic inflammation such as liver and colorectal cancers(6, 7), STAT3 activation by growth factors or interleukins suppresses T cell activation and promotes the recruitment of anti-immunity factors such as myeloid-derived suppressor cells and regulatory T cells(8, 9).

Given its fundamental roles in interpreting environmental cues to drive a cascade of signaling events that control growth and immune processes, it is essential to dissect cell type-specific roles of the JAK-STAT pathway in a pan-cancer context. This is made possible by advances in high-throughput sequencing initiatives and as many genetic alterations have become targetable, detail understanding on genetic variations would be essential to identify particular weaknesses in individual tumors in order to boost therapeutic success. We predict that genetic alterations in JAK-STAT pathway genes do not occur equally between and within cancer types. Moreover, detection of rare alterations would require a large sample size to unravel genes that are altered within specific histological subtypes of cancer. With over 18,000 samples representing 21 cancer types, we took the opportunity to systematically characterize genetic alterations within 133 JAK-STAT pathway genes to uncover shared commonalities and differences. To identify functionally relevant alterations, genetic polymorphisms were overlaid with transcript expression profiles and correlated with clinical outcomes. Our study identifies a core set of candidate JAK-STAT drivers that correlated with tumor progression and that predict overall survival outcomes in brain, renal, lung and endometrial cancers converging on similar downstream oncogenic pathways. This work provides a rich source of cancer type-dependent alterations that could serve as novel therapeutic targets to support underexploited treatment initiatives targeting JAK-STAT signaling in cancer.

## Methods

All plots were generated using R packages (pheatmap and ggplot2).

### Cancer cohorts and JAK-STAT pathway genes

133 JAK-STAT pathway genes were obtained from the Kyoto Encyclopedia of Genes and Genomes (KEGG) database listed in Table S1. Genomic, transcriptomic and clinical datasets of 21 cancer types (n=18,484) were retrieved from The Cancer Genome Atlas (TCGA)(10). The following is a list of cancer cohorts and corresponding TCGA abbreviations in parentheses: bladder urothelial carcinoma (BLCA), breast invasive carcinoma (BRCA), cervical squamous cell carcinoma and endocervical adenocarcinoma (CESC), cholangiocarcinoma (CHOL), colon adenocarcinoma (COAD), esophageal carcinoma (ESCA), glioblastoma multiforme (GBM), glioma (GBMLGG), head and neck squamous cell carcinoma (HNSC), kidney chromophobe (KICH), pan-kidney cohort (KIPAN), kidney renal clear cell carcinoma (KIRC), kidney renal papillary cell carcinoma (KIRP), liver hepatocellular carcinoma (LIHC), lung adenocarcinoma (LUAD), lung squamous cell carcinoma (LUSC), pancreatic adenocarcinoma (PAAD), sarcoma (SARC), stomach adenocarcinoma (STAD), stomach and esophageal carcinoma (STES) and uterine corpus endometrial carcinoma (UCEC).

### Copy number variation analyses

Level 4 GISTIC copy number variation datasets were downloaded from the Broad Institute GDAC Firehose(11). Discrete amplification and deletion indicators were obtained from GISTIC gene-level tables. Genes with GISTIC values of 2 were annotated as deep amplification events, while genes with values of −2 were annotated as deep (homozygous) deletion events. Shallow amplification and deletion events were annotated for genes with values of +1 and −1 respectively.

### Calculating JAK-STAT and regulatory T cell scores

A JAK-STAT 28-gene signature was developed from putative gain-or loss-of-function candidates. For each patient, 28-gene scores were calculated from the average log2 expression of signature genes: IL7, IFNG, MPL, IL11, IL2RA, IL21R, OSMR, IL20RA, IFNGR1, CDKN1A, CISH, SOCS1, IL10, IL10RA, STAT2, IL24, IL23A, PIAS3, IFNLR1, EPO, TSLP, BCL2, IL20RB, IL11RA, PTPN6, IL13, IL17D and IL15RA. Regulatory T cell (Treg) scores were determined by taking the mean expression of 31 Treg genes identified from the overlap of four Treg signatures to generate a more representative gene set that is cell type-independent(12–15).

### Multidimensional scaling, survival and differential expression analyses

We previously published detailed methods on the above analyses(16–18) and thus the methods will not be repeated here. Briefly, multidimensional scaling analyses based on Euclidean distance in Fig. 2F were performed using the vegan package in R(19). Permutational multivariate analysis of variance (PERMANOVA) was used to determine statistical significance between tumor and non-tumor samples. Survival analyses were performed using Cox proportional hazards regression and the Kaplan-Meier method coupled with the log-rank test. Predictive performance of the 28-gene signature was assessed using the receiver operating characteristic analysis. To determine the prognostic significance of a combined model uniting the JAK-STAT signature and IRF8 expression or Treg scores, patients were separated into four survival categories based on median 28-gene scores and IRF8 expression or Treg scores for Kaplan-Meier and Cox regression analyses. To determine the transcriptional effects of aberrant JAK-STAT signaling, differential expression analyses were performed on patients within the 4th versus 1st survival quartiles (stratified using the 28-gene signature).

### Functional enrichment and transcription factor analyses

Differentially expressed genes (DEGs) identified above were mapped to KEGG and Gene Ontology (GO) databases using GeneCodis(20) to determine significantly enriched pathways and biological processes. DEGs were also mapped to ENCODE and ChEA databases using Enrichr(21, 22) to identify transcription factors that were significant regulators of the DEGs.

## Results

### Copy number and transcriptome analyses reveal conserved driver mutations in JAK-STAT pathway genes

We interrogated genomic and transcriptomic landscape of 133 JAK-STAT pathway genes in 18,484 patients across 21 cancer types (Table S1). To investigate the effects of JAK-STAT pathway genomic alterations on transcriptional output, we first analyzed copy number variation (CNV) of all 133 genes. CNVs were classified into four categories: low-level amplifications, deep amplifications, low-level deletions (heterozygous deletions) and deep deletions (homozygous deletions). Lung squamous cell carcinoma (LUSC) and papillary renal cell carcinoma (KIRP) had the highest and lowest fraction of samples with deleted JAK-STAT pathway genes respectively (Table S2). In terms of gene amplification, this was also the highest in lung squamous cell carcinoma (LUSC) and the lowest in pancreatic adenocarcinoma (PAAD) (Table S2). To identify pan-cancer CNV events, we focused on genes that were deleted or amplified in at least one-third of cancer types (7 cancers). We identified 71 and 49 genes that were recurrently deleted and amplified respectively in at least 20% of samples within each cancer type and at least 7 cancer types (Table S2). Esophageal carcinoma (ESCA) had 70 genes that were recurrently deleted while only four recurrently deleted genes were found in papillary renal cell carcinoma (KIRP). When considering recurrent gene amplifications, lung adenocarcinoma (LUAD) had the highest number of gains (48 genes), while the lowest number of gene gains was observed in glioma (GBMLGG) (3 genes) (Table S2). CNV events associated with transcriptional changes could represent candidate driver genes. Loss-of-function genes can be identified from genes that were recurrently deleted and downregulated at the transcript level. Similarly, genes that were concomitantly gained and upregulated could represent a gain-of-function. Differential expression profiles (tumor vs. non-tumor) were intersected with CNV profiles and we identified 40 driver genes representing potential loss-or gain-of-function (Fig. 1). In at least 7 cancer types, 18 genes were recurrently deleted and downregulated (log2 fold-change < −0.5, P < 0.01), while a non-overlapping set of 22 genes were recurrently amplified and upregulated (log2 fold-change > 0.5, P < 0.01) (Fig. 1).

**Fig. 1.**
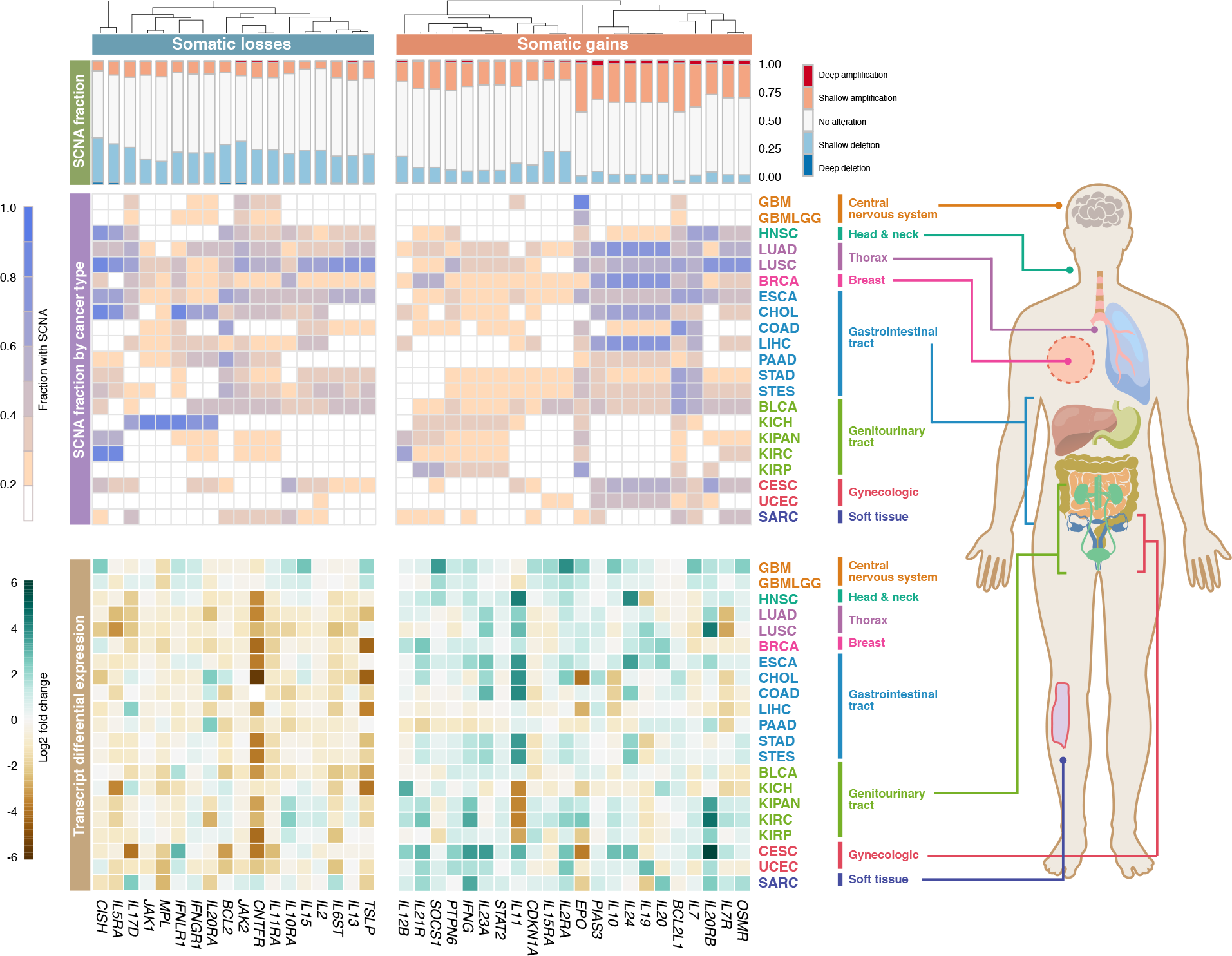
Pan-cancer genomic and transcriptomic alterations of JAK-STAT pathway genes. Stacked bar charts depict the fraction of altered samples for each of the 40 putative driver genes. Heatmap in the center illustrates the fraction of somatic gains and losses per gene within each cancer type. Heatmap below illustrates differential expression profiles (tumor vs. non-tumor) for each gene. Hierarchical clustering is performed on columns (genes) using Euclidean distance metric. Cancer abbreviations are listed in the Methods section.

### JAK-STAT driver genes predict over-all survival in diverse cancer types

Focusing on the 40 driver genes identified above, we next analyzed whether expression levels of individual genes were associated with overall survival outcomes. Cox proportional hazards regression analyses demonstrated that all 40 genes harbored prognostic information in at least one cancer type. IL11, PTPN6 and CISH were significantly associated with survival outcomes in patients from 9 cancer cohorts (Fig. 2A). In contrast, IL19, CNTFR and JAK2 were some of the least prognostic genes (Fig. 2A). When deciphering the contribution of individual genes across cancer types, we observed that the glioma (GBMLGG) cohort had the highest number of prognostic genes (31/40 genes) (Fig. 2A). On the other hand, none of the 40 driver candidates were prognostic in esophageal carcinoma (ESCA) and cholangiocarcinoma (CHOL), which suggests the minimal contribution of JAK-STAT signaling in driving tumor progression in these cancer types (Fig. 2A). To identify a core set of prognostic genes denoting pan-cancer significance, we performed Spearman’s correlation analyses on hazard ratio (HR) values obtained from Cox regression and identified 28 highly-correlated genes: IL7, IFNG, MPL, IL11, IL2RA, IL21R, OSMR, IL20RA, IFNGR1, CDKN1A, CISH, SOCS1, IL10, IL10RA, STAT2, IL24, IL23A, PIAS3, IFNLR1, EPO, TSLP, BCL2, IL20RB, IL11RA, PTPN6, IL13, IL17D and IL15RA (Fig. 2B). These genes were collectively regarded as a pan-cancer JAK-STAT signature. To determine the extent of JAK-STAT pathway variation across cancer types, we calculated an activity score based on the mean expression of the 28 genes. When cancers were sorted according to their pathway activity scores, chromophobe renal cell cancer (KICH) had the lowest median score while the highest median score was observed in head and neck cancer (HNSC) (Fig. 2C). Hierarchical clustering of the 28 driver genes demonstrated that they exhibited a wide range of expression depending on the cellular context where they could serve as potential candidates for therapy. For example, IL7, IL15RA, IL21R, IL10, OSMR, IFNGR1, IL10RA, IL2RA, IFNG, IL24, SOCS1, IL20RA, IL11 and IL23A were highly expressed in gastrointestinal cancers (PAAD, STAD and STES) (Fig. 2D). When the 28-gene scores were employed for patient stratification, we observed that the JAK-STAT signature conferred prognostic information in five diverse cancer cohorts (Fig. 2E). Intriguingly, the significance of the signature in predicting overall survival was cancer type-dependent. Kaplan-Meier analyses and log-rank tests revealed that patients with high scores (4th quartile) had higher death risks in glioma (P<0.0001), pan-kidney (consisting of chromophobe renal cell, clear cell renal cell and papillary renal cell cancers; P<0.0001) and clear cell renal cell (P<0.0001) cohorts (Fig. 2E). In contrast, high expression of signature genes was linked to improved survival rates in lung (P=0.025) and endometrial (P=0.032) cancers (Fig. 2E). These results were independently corroborated using Cox regression analyses: glioma (HR=6.832, P<0.0001), pan-kidney (HR=3.335, P<0.0001), clear cell renal cell (HR=4.292, P<0.0001), lung (HR=0.624, P=0.028) and endometrium (HR=0.504, P=0.027) (Table S3). Since high expression of signature genes was associated with poor survival outcomes in brain and renal cancers, as expected, we observed a significant increase in expression scores according to tumor stage (Fig. 2G). An opposite trend was observed in lung and endometrial cancers where more aggressive tumors had lower expression scores (Fig. 2F). Lastly, multidimensional scaling analyses of signature genes in the five cohorts revealed significant differences between tumor and non-tumor samples, implying that dysregulated JAK-STAT signaling may serve as a diagnostic marker for early detection in pre-cancerous lesions (Fig. 2F).

**Fig. 2.**
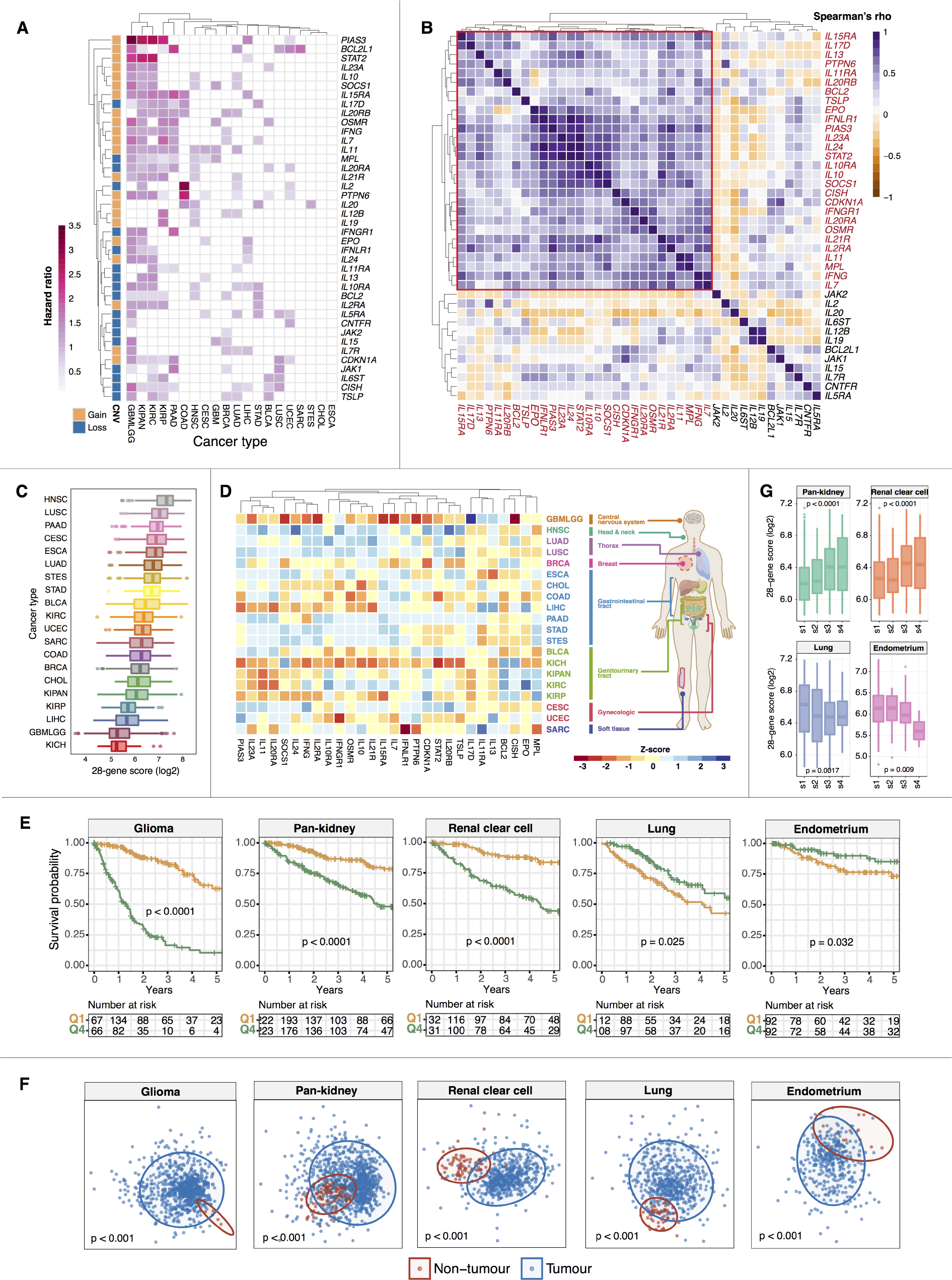
Prognostic significance of JAK-STAT driver genes. (A) Heatmap illustrates hazard ratio values obtained from Cox proportional hazards regression on 40 candidate drivers across all cancer types. (B) Heatmap illustrates Spearman’s correlation coefficient values comparing hazard ratios of the 40 driver genes. Highly correlated genes are highlighted in red and are demarcated by a red box. (C) Box plots represent the distribution of 28-gene scores derived from highly correlated JAK-STAT driver genes. Cancers are ranked from high to low median scores. (D) Heatmap depicts the Z-scores for each of the 28 genes by cancer types. (E) Kaplan-Meier analyses confirmed prognosis of the 28-gene signature in five cancer cohorts. Patients are separated into 4th and 1st survival quartiles based on their 28-gene scores. P values are obtained from log-rank tests. (F) Ordination plots of multidimensional scaling analyses using signature genes reveal significant differences between tumor and non-tumor samples. P values are obtained from PERMANOVA tests. (G) Expression distribution of 28-gene scores in tumors stratified by stage (s1, s2, s3 and s4). P values are determined using ANOVA.

### The JAK-STAT 28-gene signature is an independent prognostic factor

Multivariate Cox regression analyses confirmed that the signature was independent of tumor, node and metastasis (TNM) stage: glioma (HR=2.377, P=0.018), pan-kidney (HR=2.468, P<0.0001), clear cell renal cell (HR=2.552, P=0.00047), lung (HR=0.636, P=0.031) and endometrium (HR=0.434, P=0.033) (Table S3). Given that the signature was an independent predictor of overall survival, we reasoned that its predictive performance could be increased when used in conjunction with TNM staging. Employing the receiver operating characteristic (ROC) analysis, we demonstrated that a combined model uniting the signature and TNM staging could outperform the signature (higher area under the curve [AUC] values) when it was considered alone: pan-kidney (0.838 vs. 0.800), clear cell renal cell (0.836 vs. 0.789), lung (0.724 vs. 0.703) and endometrium (0.760 vs. 0.713) (Fig. 3A). Independently, Kaplan-Meier analyses and log-rank tests confirmed that the signature allowed further delineation of risk groups within similarly-staged tumors: pan-kidney (P<0.0001), clear cell renal cell (P<0.0001), lung (P<0.0001) and endometrium (P<0.0001) (Fig. 3B).

**Fig. 3.**
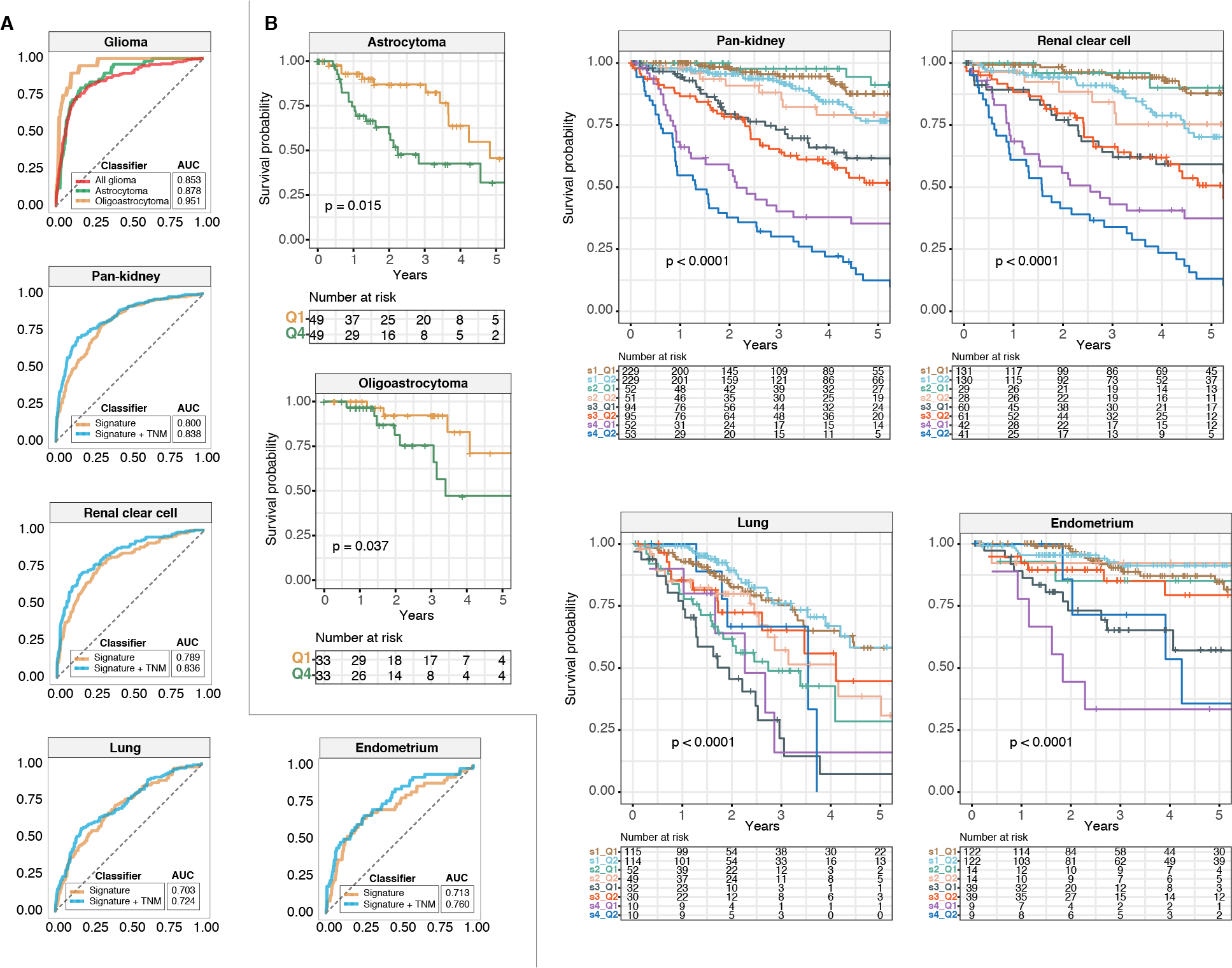
The 28-gene signature is independent of tumor, node and metastasis (TNM) stage. (A) Receiver operating characteristic analysis is used to assess the predictive performance of the signature and TNM stage. For glioma patients, area under the curves (AUCs) are compared between histological subtypes. (B) Kaplan-Meier analyses of patients stratified by tumor stage or, in the case of glioma, by histological subtype and the 28-gene signature. For histological subtypes of glioma, log-rank tests are used to compare patients within the 1st and 4th survival quartiles. For the other cancers, patients are first stratified according to TNM stage followed by median-stratification into low-and high-score groups using the 28-gene signature. P values are obtained from log-rank tests.

High JAK-STAT scores were associated with decreased survival rates in glioma patients. We confirmed that this was also true for histological subtypes of glioma: astrocytoma (P=0.015) and oligoastrocytoma (P=0.037) (Fig. 3B). Independently confirmed using Cox regression, patients within the 4th survival quartile had lower survival rates: astrocytoma (HR=2.377, P=0.018) and oligoastrocytoma (HR=2.730, P=0.038) (Table S3). In terms of the signature’s predictive performance, ROC analyses revealed that it performed the best in oligoastrocytoma (AUC=0.951), followed by astrocytoma (AUC=0.878) and glioma patients when considered as a full cohort (AUC=0.853) (Fig. 3A).

### Consequences of dysregulated JAK-STAT signaling and significant crosstalk with tumor immunity

Since dysregulated JAK-STAT signaling was associated with survival outcomes (Fig. 2 and Fig. 3), we reasoned that patients from diverse cancer types might harbor similar transcriptional defects caused by aberrant activation of JAK-STAT. Differential expression analyses performed between the 4th and 1st quartile patients revealed that a striking number of over 200 differentially expressed genes (DEGs) were shared between all five prognostic cohorts (Fig. 4A; Table S4). Significant overlaps were observed in DEGs where 555 genes were found in at least four cohorts, 1,009 genes in at least three cohorts and 2,034 genes in at least two cohorts (Fig. 4A; Table S4). The highest number of DEGs was observed in glioma (2,847 genes), followed by pan-kidney (2,810 genes), lung (1,221 genes), clear cell renal cell (1,161 genes) and endometrial (782 genes) cancers (Fig. 4A; Table S4). To determine their functional roles, the DEGs were mapped to Gene Ontology (GO) and KEGG databases. All five cohorts exhibited remarkably similar patterns of enriched biological processes (Fig. 4B). Pan-cancer enrichments of ontologies related to inflammation and immune function were observed, e.g., cytokine and chemokine signaling, T cell and B cell receptor signaling, natural killer cell-mediated processes, Toll-like receptor signaling and NOD-like receptor signaling (Fig. 4B). Additionally, genes associated with other oncogenic pathways (Wnt, MAPK, TGF-β, PPAR and VEGF)(23–25) were frequently altered at both transcriptional and genomic levels (Fig. 4B and 4C). CNV analyses performed on DEGs that were found to be in common in at least three cohorts demonstrated that transcriptional dysregulation of the aforementioned oncogenic pathways was attributed to activating or inactivating CNVs (Fig. 4C). For instance, except for THBS1 and THBS2, a vast majority of TGF-β DEGs exhibited somatic gains (Fig. 4C). To further explore which upstream transcriptional regulators were involved, we mapped the DEGs to ENCODE and ChEA databases (Fig. 4B). Interestingly, we observed significant enrichment of transcription factors (TFs) involved in modulating immune function: IRF8(26), RUNX1(27), RELA(28) and EZH2(29) (Fig. 4B). Taken together, our analyses revealed that pan-cancer JAK-STAT drivers underpin numerous aspects of tumor oncogenesis and immunity, which play important roles in tumor progression and ultimately patient prognosis.

**Fig. 4.**
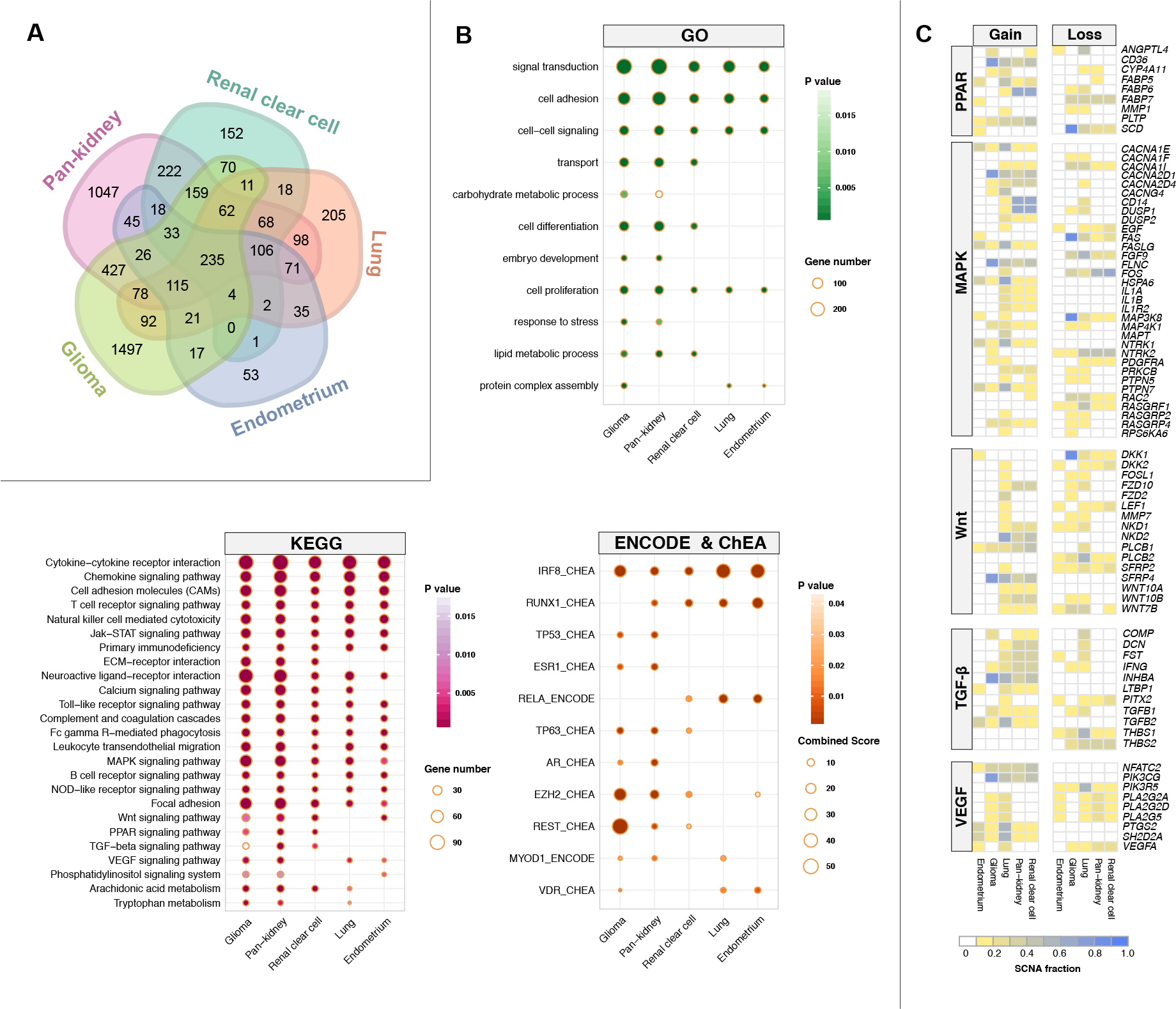
Aberrant JAK-STAT signaling drives malignant progression through crosstalk with other oncogenic pathways. (A) Venn diagram depicts the number of overlapping differentially expressed genes (DEGs) in five cohorts. Differential expression analyses are performed between patients stratified into the 4th and 1st survival quartiles using the JAK-STAT signature in five cancer cohorts. (B) Functional enrichment analyses are performed by mapping DEGs to the Gene Ontology and KEGG databases. Mapping of DEGs to ENCODE and ChEA databases identify enriched transcription factor binding associated with DEGs. (C) Somatic copy number alteration frequencies for DEGs identified from enriched pathways in B. Heatmaps depict the fraction of somatic gains and losses for DEGs found within five oncogenic pathways. Only genes that are altered in more than 10% of samples within each tumor type are shown.

### IRF8 and JAK-STAT pathway synergistically influence survival outcomes in glioma and renal cancer

IRF8 was among one of the most enriched TFs implicated in the regulation of transcriptional outputs of patients with dysregulated JAK-STAT signaling (Fig. 4B). As a member of the interferon regulatory factor family, IRF8 is needed for the development of immune cells and is often regarded as a tumor suppressor gene since a loss of function is associated with increased metastatic potential(30, 31). Thus, we predict that tumors with low expression of IRF8 would be more aggressive. To evaluate the combined relationship between JAK-STAT signaling and IRF8 expression, patients were categorized into four groups based on median IRF8 and JAK-STAT scores: 1) high-high, 2) low-low, 3) high IRF8 and low 28-gene score and 4) low IRF8 and high 28-gene score. The combined model encompassing JAK-STAT and IRF8 offered an additional resolution in patient stratification: glioma (full-cohort, P<0.0001), astrocytoma (P=0.0007), pan-kidney (P<0.0001) and clear cell renal cell (P<0.0001) (Fig. 5A). Indeed, patients with low IRF8 and high 28-gene scores performed the worst in cancers where hyperactive JAK-STAT signaling was linked to adverse survival outcomes: glioma (full-cohort: HR=5.826, P<0.0001), astrocytoma (HR=3.424, P=0.0032), pan-kidney (HR=5.131, P<0.0001) and clear cell renal cell (HR=5.389, P<0.0001) (Fig. 5B). Our results support a model in which IRF8 influences the behavior of tumors with aberrant JAK-STAT signaling.

**Fig. 5.**
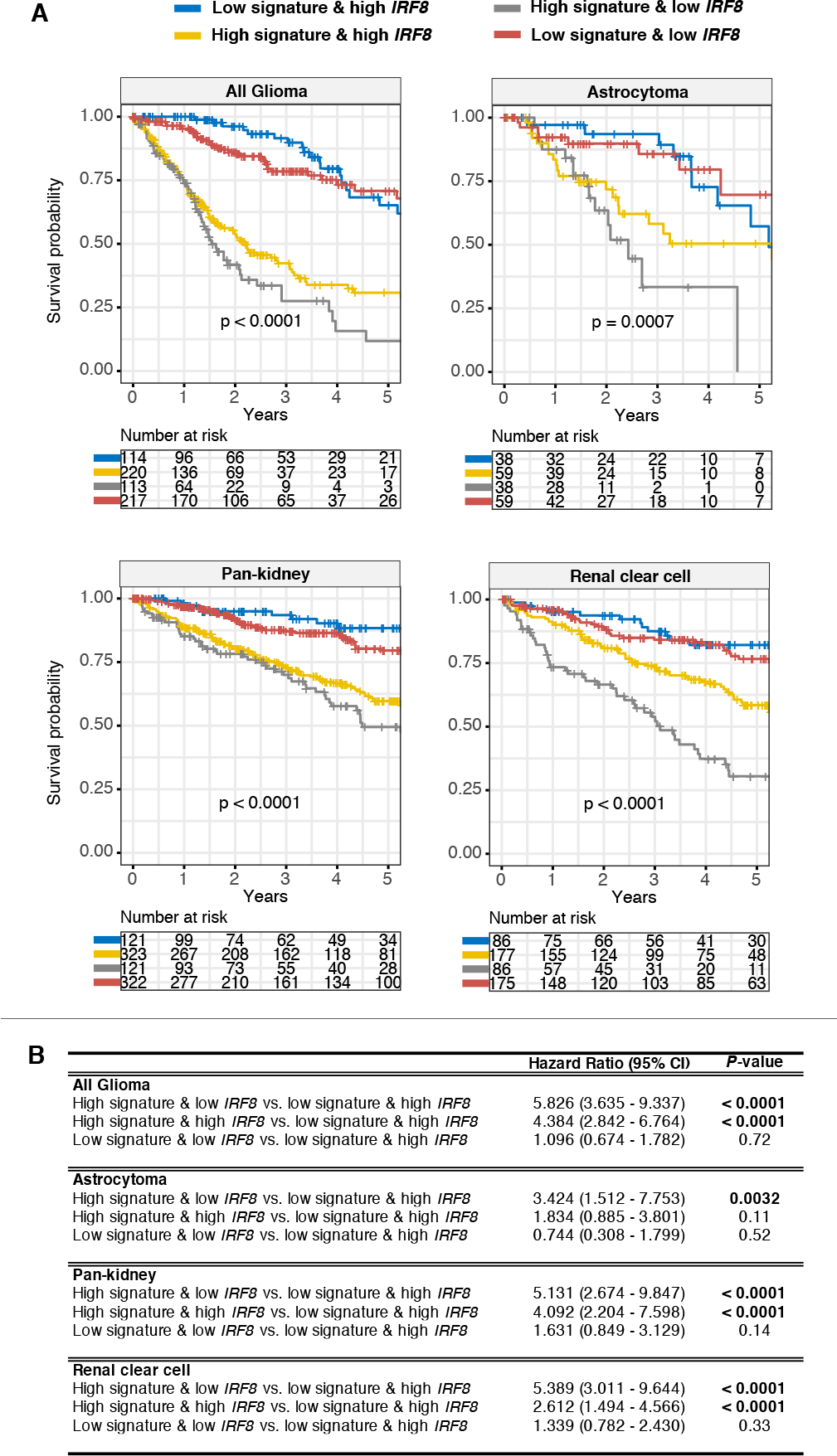
Prognostic relevance of the crosstalk between JAK-STAT signaling and IRF8. (A) Patients are grouped into four categories based on median 28-gene and IRF8 expression values. Kaplan-Meier analyses are performed on the four patient groups to determine the ability of the combined JAK-STAT-IRF8 model in determining over-all survival in glioma and renal cancers. P values are obtained from log-rank tests. (B) Table inset shows univariate Cox proportional hazards analyses of the relation-ship between JAK-STAT signaling and IRF8. Significant P values are highlighted in bold. CI = confidence interval.

### Hyperactive JAK-STAT signaling attenuates tumor immunity

Given the wide-ranging effects of JAK-STAT signaling on immune-related functions, we hypothesized that JAK-STAT activity would correlate with immune cell infiltration. We retrieved genes implicated in regulatory T cell (Treg) function from four studies and isolated 31 genes that were common in all four Treg signatures(12–15). Treg scores were calculated for each patient based on the mean expression levels of the 31 genes. Remarkably, we observed strong positive correlations between Treg and JAK-STAT scores, suggesting that tumor tolerance was enhanced in patients with hyper-active JAK-STAT signaling: glioma (rho=0.85, P<0.0001), pan-kidney (rho=0.88, P<0.0001) and clear cell renal cell (rho=0.83, P<0.0001) (Fig. 6A). As in the previous section, patients were stratified into four categories based on median JAK-STAT and Treg scores for survival analyses. Log-rank tests and Cox regression analyses confirmed that elevated Treg activity further exacerbated disease phenotypes in patients where JAK-STAT scores were already high: glioma (full cohort, HR=6.183, P<0.0001), astrocytoma (HR=3.035, P=0.00042), pan-kidney (HR=3.133, P<0.0001) and clear cell renal cell (HR=2.982, P<0.0001) (Fig. 6B and 6C). Taken together, elevated JAK-STAT signaling may increase the ability of tumors to escape immunosurveillance, resulting in more aggressive tumors and increased mortality rates.

**Fig. 6.**
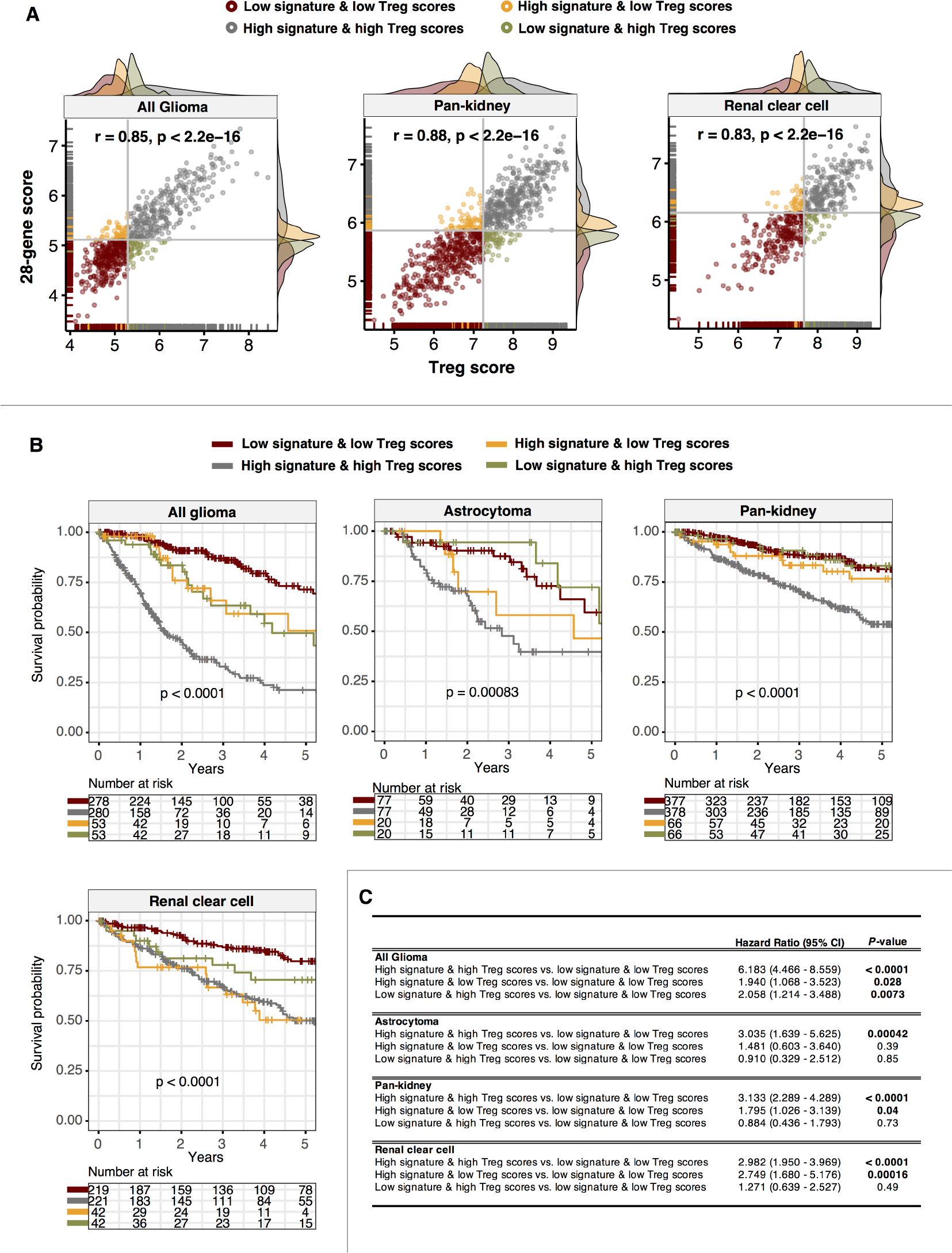
Tumors with hyperactive JAK-STAT signaling are hypoimmunogenic. (A) Scatter plots depict significant positive correlations between 28-gene and regulatory T cell (Treg) scores in glioma and renal cancer patients. Patients are grouped into four categories based on median 28-gene and Treg scores. Density plots at the x-and y-axes show the distribution of 28-gene and Treg scores. (B) Kaplan-Meier analyses are performed on the four patient groups to determine the ability of the combined JAK-STAT-Treg model in determining overall survival in glioma histological subtypes and renal cancer. P values are obtained from log-rank tests. (C) Table inset shows univariate Cox proportional hazards analyses of the relationship between JAK-STAT signaling and anti-tumor immunity. Significant P values are highlighted in bold. CI = confidence interval.

## Discussion

Tumor-promoting and tumor-suppressing roles of JAK-STAT signaling is very much cell type-dependent (Fig. 1, Fig. 2, Fig. 3). JAK-STAT activation appears to drive oncogenic progression in liver cancer(32) and infection with hepatitis viruses could also induce pathway activation(33, 34). Moreover, liver tumors with downregulated SOCS proteins (inhibitors of JAK-STAT signaling) are associated with poor prognosis(32). JAK and STAT3 promote cell proliferation, invasion and migration in colorectal cancer through the regulation of cell adhesion molecules and growth factors(35). In contrast, phosphorylated STAT5 promotes cellular differentiation and inhibits invasive properties in breast cancer cells(36, 37). A decrease in STAT5 is also linked to poorly differentiated morphology and advance histological grades in breast tumors(38, 39). Similarly, in rectal cancers, patients with tumors positive for phosphorylated STAT3 had improved survival outcomes(40). In contrast, high levels of phosphorylated STAT3 is associated with reduced survival rates in glioblastoma(41) and renal cancer(42), which independently corroborates our findings on the tumor-promoting effects of JAK-STAT signaling in these cancer types (Fig. 2 and 3). Given its ambiguous role, understanding the function of JAK-STAT signaling in a pan-cancer context would be necessary to increase the success of therapy in tumors with abnormal pathway activity. In an integrated approach employing genomic, transcriptomic and clinical datasets, we elucidated pan-cancer patterns of JAK-STAT signaling converging on a core set of candidate driver genes known as the JAK-STAT 28-gene signature (Fig. 2). We demonstrated prognosis of the signature in five cancer cohorts (n=2,976), where its performance was independent of TNM stage (Fig. 2 and 3). Patients with aberrant JAK-STAT signaling exhibited interactions with other major oncogenic pathways, including MAPK, Wnt, TGF-β, PPAR and VEGF (Fig. 4). This suggests that co-regulation of intracellular signaling cascades could have direct functional effects and combinatorial therapies simultaneously targeting these pathways may improve treatment efficacy and overcome resistance.

We demonstrated that hyperactivation of JAK-STAT signaling promotes the loss of anti-tumor immunity in glioma and renal cancer patients (Fig. 6). In tumor cells, constitutive activation of STAT3 inhibits anti-tumor immune response by blocking the secretion of proinflammatory cytokines and suppressing dendritic cell function(43). Furthermore, hyperactivation of STAT3 is linked to abnormal differentiation of dendritic cells in colon cancer cells(44). STAT3 promotes interleukin-10-dependent Treg function(45) while STAT5 promotes Treg differentiation(46). We demonstrated that in glioma and renal cancer, JAK-STAT scores were strongly correlated with Treg expression scores, suggesting that persistent activation of JAK-STAT could promote tumor immune evasion in these cancers (Fig. 6). Moreover, in tumors with high levels of JAK-STAT signaling and low IRF8 expression (a TF involved in regulating innate and adaptive immune responses), we observed a dramatic decrease in overall survival rates (Fig. 5). IRF8 is essential for dendritic cell development and proatherogenic immune responses(47). Moreover, IRF8 is crucial for NK-cell-mediated immunity against mouse cytomegalovirus infection(48). Promoter hypermethylation and gene silencing of IRF8 abrogates cellular response to interferon stimulation and overexpression of IRF8 in nasopharyngeal, esophageal and colon cancer cell lines could inhibit clonogenicity(31). IRF8 expression is also negatively correlated with metastatic potential by increasing tumor resistance to Fas-mediated apoptosis(30). Together, our results and those of others support the tumor suppressive roles of IRF8. Importantly, loss of IRF8 may further suppress tumor immunity in patients with hyperactive JAK-STAT signaling. A number of JAK inhibitors (tofacitinib, ruxolitinib and oclacitinib) have been FDA-approved and along with 2nd-generation JAKinibs and STAT inhibitors currently undergoing testing(49), our signature can be used for patient stratification before adjuvant treatment with these inhibitors to enable selective targeting of tumors that are likely to respond.

## Conclusion

What started as an initiative to understand JAK-STAT pathway genes that are somatically altered in varying combinations and frequencies across diverse cancer types has now resulted in a framework that supports selective targeting of novel candidates in a broad spectrum of cancers. Our study also reveals important crosstalk between JAK-STAT and other oncogenic pathways and when targeted together, this could radically improve clinical outcomes. Other researchers can harness this novel set of data and the JAK-STAT gene signature in the future for prospective validations in clinical trials and functional studies involving JAK-STAT inhibitors and immune checkpoint blockade.

